# Constrained mutational sampling of amino acids in HIV-1 protease evolution

**DOI:** 10.1101/354597

**Authors:** Jeffrey I. Boucher, Troy W. Whitfield, Ann Dauphin, Gily Nachum, Carl Hollins, Konstantin B. Zeldovich, Ronald Swanstrom, Celia A. Schiffer, Jeremy Luban, Daniel N. A. Bolon

## Abstract

The evolution of HIV-1 protein sequences should be governed by a combination of factors including nucleotide mutational probabilities, the genetic code, and fitness. The impact of these factors on protein sequence evolution are interdependent, making it challenging to infer the individual contribution of each factor from phylogenetic analyses alone. We investigated the protein sequence evolution of HIV-1 by determining an experimental fitness landscape of all individual amino acid changes in protease. We compared our experimental results to the frequency of protease variants in a publicly available dataset of 32,163 sequenced isolates from drug-naïve individuals. The most common amino acids in sequenced isolates supported robust experimental fitness, indicating that the experimental fitness landscape captured key features of selection acting on protease during viral infections of hosts. Amino acid changes requiring multiple mutations from the likely ancestor were slightly less likely to support robust experimental fitness than single mutations, consistent with the genetic code favoring chemically conservative amino acid changes. Amino acids that were common in sequenced isolates were predominantly accessible by single mutations from the likely protease ancestor. Multiple mutations commonly observed in isolates were accessible by mutational walks with highly fit single mutation intermediates. Our results indicate that the prevalence of multiple base mutations in HIV-1 protease is strongly influenced by mutational sampling.

## Introduction

Mutations are important to HIV-1 and many other viruses because they enable evasion of immune recognition (Nowak, Anderson et al. 1991) and escape from anti-viral drug treatments (Kantor, Shafer et al. 2004). For example, when the mutation rate was reduced by engineering a high-fidelity polymerase in polio, the resulting viruses had an impaired ability to infect and colonize in mice (Pfeiffer and Kirkegaard 2005; Vignuzzi, Stone et al. 2006). Similar observations of reduced infectivity for engineered high-fidelity viruses have been made in chikungunya virus (Coffey, Beeharry et al. 2011) and human enterovirus 71 (Meng and Kwang 2014). In HIV-1, mutations that either increase or decrease the mutation rate show reduced replication efficiency in cell culture (Dapp, Heineman et al. 2013), suggesting that the mutation rate of HIV-1 has been subject to natural selection.

HIV-1 generates mutations as an inherent part of its infection cycle. HIV-1 is an RNA virus that replicates in host cells through a DNA intermediate. The process of reverse transcribing viral RNA into DNA is error prone and is a main contributor to the mutations that HIV-1 accumulates (Preston, Poiesz et al. 1988; Cuevas, Geller et al. 2015). The error rate of HIV-1 replication in cell culture has been measured as 3 × 10^−5^ mutations per base per replication cycle (Mansky 1996). This corresponds to 1 mutation for every 2-3 genomes of HIV-1. In addition to errors that occur during genome copying, additional mutations are caused after synthesis by host cytidine deaminases. Recent analyses of the frequency of null alleles of HIV-1 integrated as proviruses within host DNA, but incapable of infection (Cuevas, Geller et al. 2015), indicate that the *in vivo* error rate may be higher than estimates from cell culture. Of note, these findings may be influenced by selection pressure that should disfavor functional proviruses that would kill the parental cell unless silenced. Because of the large number of virions in an infected individual, even conservative estimates of the error rate lead to tremendous genetic diversity. In a widely cited review (Perelson 2002), modeling of HIV-1 infection kinetics indicated that, “mutations will occur in every position in the genome multiple times each day and that a sizeable fraction of all possible double mutations will also occur.”

While mutations in HIV-1 occur at the nucleic acid level, many of the functional impacts are caused by changes in the encoded protein sequence (Zanini, Puller et al. 2017). The amino acid changes that are efficiently sampled by HIV-1 depend on multiple factors including the probabilities of different types of nucleic acid mutations and the genetic code. Single nucleotide mutations are the most frequent errors generated when HIV-1 replicates (Mansky 1996; Abram, Ferris et al. 2010). Amino acids that are accessible by a single mutation from the parental sequence should be more frequently sampled during HIV-1 infection and evolution. Within the genetic code, amino acids with similar physical and chemical properties are encoded by nearby codons making it likely that single nucleotide mutations will cause conservative amino acid changes (Woese 1965). Detailed analyses indicate that strong selection acts on the genetic code (Sengupta and Higgs 2015) and that only one in a million random genetic codes is as effective at minimizing disruptive amino acid changes from single base mutations (Freeland and Hurst 1998). Amino acids changes that require multiple-base mutations could be infrequent due in part to disruptive amino acids causing fitness defects.

Here, we sought to understand how the genetic code impacts protein evolution in HIV-1 where most if not all single nucleotide mutations are sampled within an infected individual. We chose to focus on protease for a number of reasons including a wealth of available sequence data, the relatively small size of the 99 amino acid gene that expedited experimental analyses, and the lack of antibody pressure on protease during infection that facilitated comparison of cell culture experiments with selection in hosts. Of note, mutations in protease can influence immune recognition of peptides displayed on major histocompatibility complex of infected cells, but it is not clear how this impacts selection on protease sequence (Mueller, Spriewald et al. 2011). We observed that many amino acids requiring multiple mutations were infrequent in sequenced isolates yet were compatible with efficient viral replication. Despite the extensive genetic diversity of HIV-1 within an infected individual, our results indicate that inefficient sampling of amino acids requiring multiple mutations in protease has strongly impacted it’s evolution.

## Results and Discussion

### Analyses of protease sequences from isolates

To explore mutation and selection acting on circulating HIV-1 in the absence of drug pressure, we analyzed the amino acid sequence of protease (Shafer, Jung et al. 2000) from 30,987 protease-inhibitor-naïve people infected with subtype B HIV-1 (Fig. 1). We calculated the frequency of each amino acid at each position in protease as an estimate of fitness independent of genetic background. As with many previous analyses of HIV-1 sequence evolution (Humphris-Narayanan, Akiva et al. 2012; Ferguson, Mann et al. 2013; Butler, Barton et al. 2016; Flynn, Haldane et al. 2017; Louie, Kaczorowski et al. 2018), we assume that the sequence prevalence in isolates represent a steady state distribution. Evolutionary simulations have shown that immune pressure on HIV-1 effectively performs extensive sampling of viral sequence space, making the steady-state assumption reasonable (Shekhar, Ruberman et al. 2013). In addition, fitness landscapes of HIV-1 inferred from multiple-sequence alignments directly, with no correction for ancestry, correlate with experimental fitness measurements (Ferguson, Mann et al. 2013). The steady-state assumption is counter-intuitive because evolution is dynamic by nature, and while there is abundant evidence showing that protease sequence is currently at steady-state, there is no guarantee that this will remain the case indefinitely. Of note, the steady-state assumption is also consistent with the experimental results in this work when variation in experimental measurement are taken into consideration.

**Figure 1.**
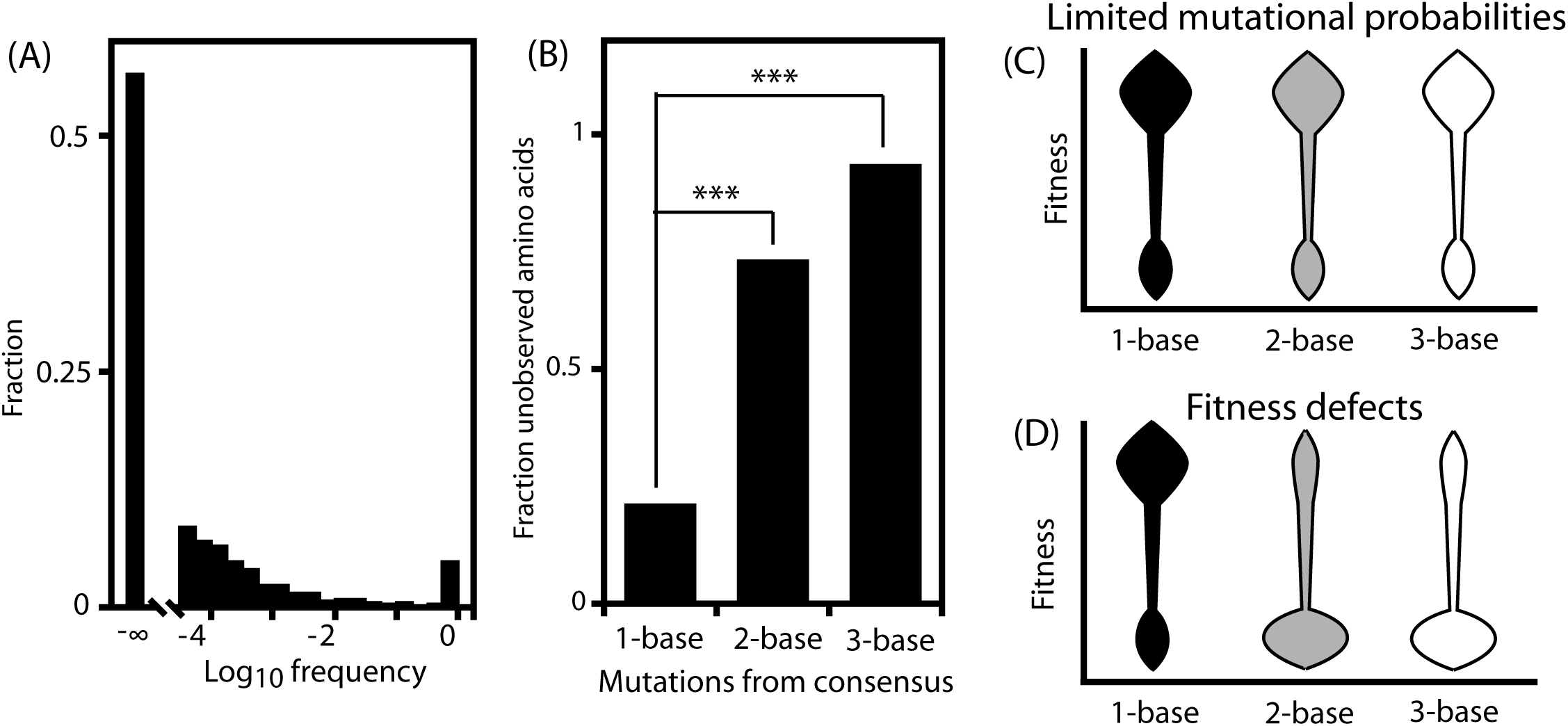
Amino acid frequencies of patient derived HIV-1 protease indicate biased accumulation of single-base mutations. (A) Frequency distribution of amino acids observed in sequenced isolates. More than 50% of amino acids were not observed. (B) The fraction of unobserved amino acids requiring 1-, 2-, or 3-mutations from a likely ancestral protein sequence. We used a chi-square test for statistical significance and *** indicates p<0.00001. (C) Depiction of fitness landscapes where all mutations exhibit similar fitness and biases for single mutations would come entirely from mutational probabilities (D) Potential landscapes that exhibit strong fitness preference for amino acid changes accessible by single nucleotide conversions.

Analyses of HIV-1 sequences indicate a symmetric radial or star pattern of evolution (Flynn, Haldane et al. 2017), such that the consensus sequence is a likely ancestral sequence. Consistent with this line of reasoning, we observed that the consensus sequence is the most abundant sequence, representing 2.5% of sequenced isolates (Fig. S1). The sequence of protease in our experimental system (pNL4.3) is the consensus of the sequenced isolates indicating that it represents the likely ancestral sequence.

The majority of possible amino acids were observed rarely or not at all in isolates (Fig. 1A), indicating that protease function is subject to strong purifying selection and that the likely ancestral sequence is highly adapted. Amino acids that were not accessible from the consensus by single mutations were under-represented in the sequenced isolates. While 79% of amino acids that were one mutation from the consensus were observed in isolates, 73% of amino acids two base substitutions away were not observed (Fig. 1B). There are two main reasons that an amino acid may be at low frequency in a population: mutation and selection. If an amino acid causes a fitness defect, selection will drive it to low frequency. For neutral amino acids without any fitness effects, the probability of the amino acid will be proportional to the rate at which mutations cause the amino acid (Kimura 1983). The disproportionate fraction of multiple mutations that were unobserved could be due to limited mutational probabilities (Fig. 1C), and/or fitness distributions skewed towards deleterious relative to single mutations (Fig. 1D). We investigated these possibilities further by experimentally determining a protein fitness landscape for HIV-1 protease.

### Quantification of experimental fitness

We performed an EMPIRIC (Hietpas, Jensen et al. 2011) mutational scan of the HIV-1 protease gene as outlined in Figure 2A. We individually randomized each amino acid position of protease in the NL4-3 strain of HIV-1 and quantified the experimental fitness of each possible mutation during infection of a T-cell line using a deep sequencing readout that our lab had previously developed and adapted to analyze HIV-1 (Duenas-Decamp, Jiang et al. 2016). Based on our previous work, the errors from processing and sequencing using this approach are a small fraction compared to the frequency of the vast majority of mutations in our initial libraries. Stop mutations provide an additional useful check on the reliability of the experimental measurements. For example, if the frequency of a stop mutation were primarily due to sequencing errors, then it may persist in the sequencing readout and appear highly fit. Fitness measurements were scaled as selection coefficients (s) where 0 corresponds to wildtype (WT) or neutral and −1 corresponds to a null allele defined based on premature stop codons (see Methods). Across the sequence of protease, premature stop codons cause severe fitness defects (Table S1), indicating that sequencing noise rarely if ever prevents observation of strong fitness defects.

**Figure 2.**
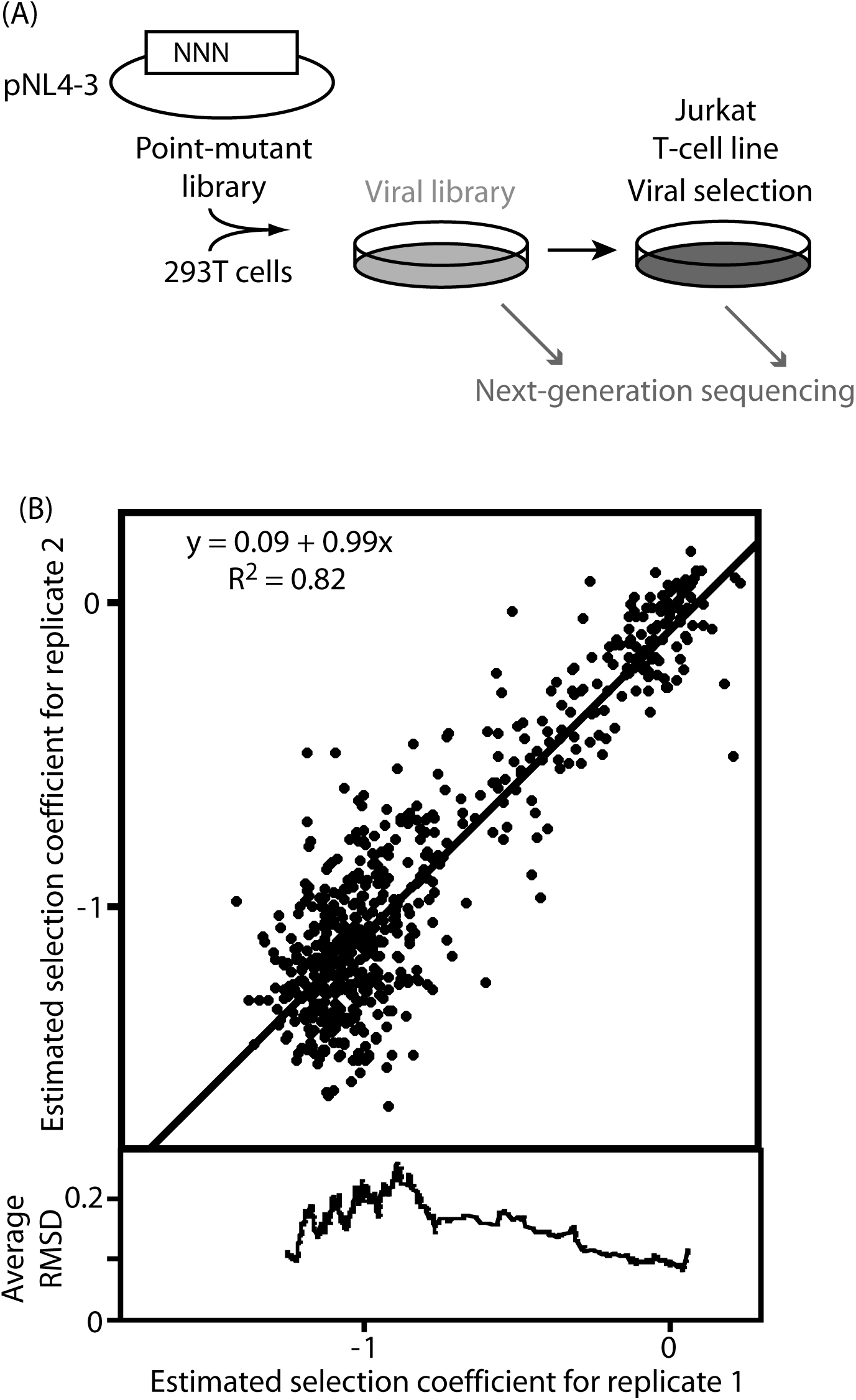
EMPIRIC measurement of selection coefficients. (A) Point mutations were generated in pNL4-3 and transfected into 293T cells to generate viral libraries that were expanded in Jurkat cells. Next-generation sequencing was used to quantify the change in frequency of mutations in the viral library before, during and after passage in Jurkat cells. (B) Comparison of estimated selection coefficients for amino acid changes from experimental replicates for three regions of protease (amino acids 20-29, 50-59, 80-89). The bottom panel shows RMSD averaged for a running window of 40 data points.

Synonymous mutations provide an additional type of internal control. Across the sequence of protease, synonymous mutations tended to exhibit estimated selection coefficients centered on 0 (Fig. S2), suggesting that fitness impacts were primarily determined by amino acid changes. Of note, synonymous mutations showed statistically significant deviations from neutral expectations in two 14 amino acid windows: at the beginning of protease and centered at amino acid position 54 (Fig. S2). The estimated selection coefficients of synonymous mutations at the beginning of protease are consistent with selection on the amino acid sequence of the p6 reading frame that overlaps with this region of protease (Fig. S3) as has been previously observed for the region of overlap between tat and rev genes in HIV-1 (Fernandes, Faust et al. 2016). We do not have explanations for the observed effects of synonymous mutations at the region centered on position 54 of protease. The effects of synonymous mutations between experimental replicates showed weak correlation (R^2^=0.01, Fig. S2), suggesting that much of the observed effects arose from experimental variation or noise. To reduce the impacts of noise, we chose to estimate the effects of each amino acid change in protease by summing sequencing counts over all synonyms (Table S1).

To assess the reliability of our fitness estimates, we compared estimated selection coefficients for technical repeats for three regions encompassing 30 of the 99 amino acid positions in protease (Fig. 2B). The estimated selection coefficients for amino acids tended to cluster around neutral (s=0) and null (s=-1), as has been almost universally observed for systematic or random mutation studies (Jiang, Mishra et al. 2013; Canale, Cote-Hammarlof et al. 2018). The neutral cluster and the null cluster are well distinguished in both experimental replicates. Overall the experimental replicates were linearly correlated with an intercept close to 0, a slope close to 1, and R^2^=0.82 indicating that normalization of data between replicates was effective. Analyses of the difference between replicate experiments indicates that small effect mutations had less measurement variance than mutations with severe fitness defects (Fig. 2B) consistent with greater sampling of fit variants in sequencing.

### Single and double nucleotide mutations

Single base mutations tend to cause chemically conservative amino acid changes compared to changes that are more distant in the genetic code, requiring two or more mutations. We examined the experimental protein fitness landscape accessible by single and multiple mutations (Fig. 3). To account for the potential accumulation of synonymous mutations, we considered the minimum number of mutations required to change between each pair of amino acids. Using this approach there were 827 single mutations from the likely ancestral protease sequence, and 1063 double mutations. Because the third position in codons is often degenerate, there are very few amino acid changes the require triple mutations, and there were 90 for protease. Because of the small number of triple mutations and biases in where they were located on the protease structure, we focused our analyses on single and double mutations.

**Figure 3.**
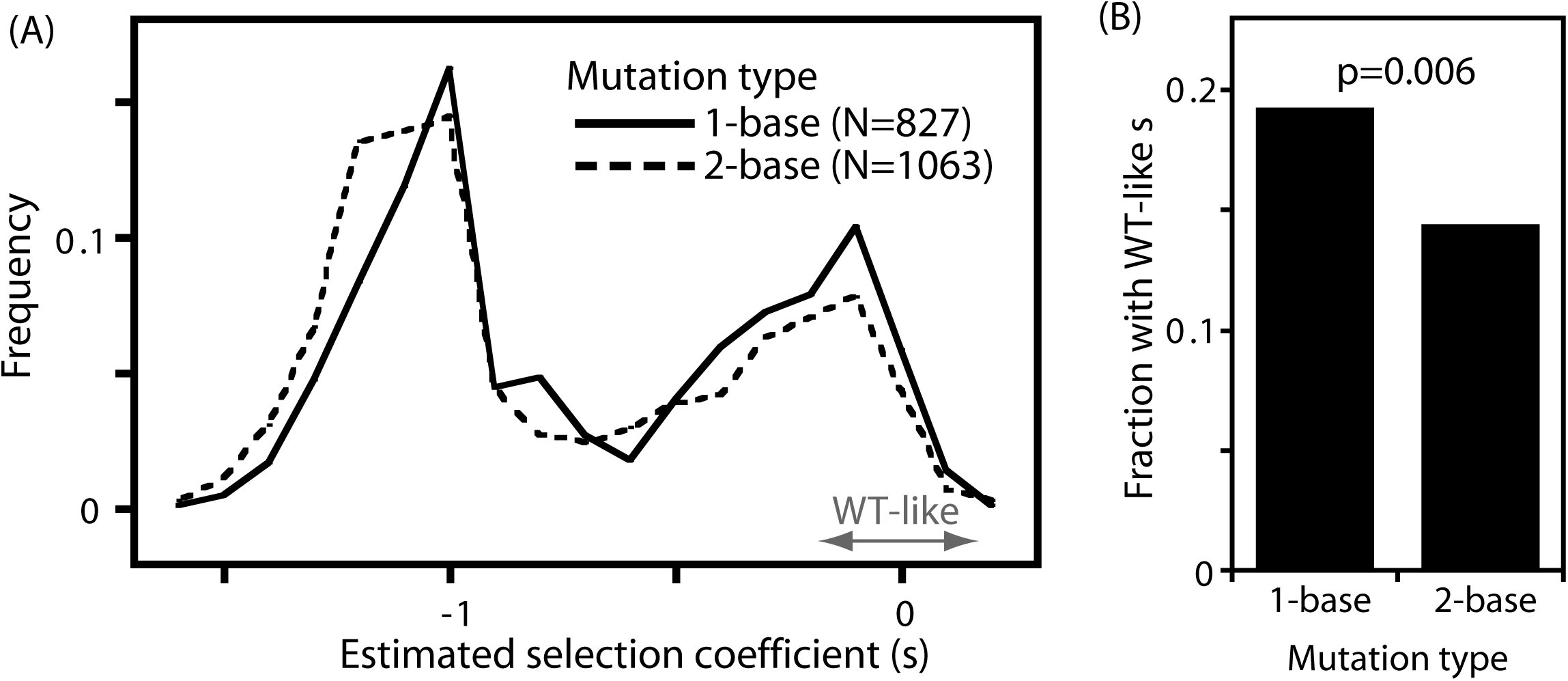
Distribution of estimated selection coefficients for amino acid changes that require 1-, 2-, or 3-nucleotide mutations. The 1-base and 2-base distributions are different (p=10-6) based on a two-sample Kolmogorov-Smirnov test. The range of WT-like selection coefficients was defined as the twice the standard deviation of WT synonyms. (B) The fraction with WT-like fitness are different for 1-base and 2-base mutations. Statistical significance was determined using a chi-square test.

The distributions of estimated selection coefficients for both single and double mutations were bi-modal with clusters near null and neutral (Fig. 3A). Indicating that the tendency for amino acid changes to cause all or none impacts on fitness is not dependent on single or double mutations. This observation is consistent with the capability of both single and double mutations to cause small or large changes in the physical properties of amino acids. For example, a single mutation can cause a small physical change from aspartate to glutamate, or a large physical change from aspartate to tyrosine. The genetic code biases for conservative amino acid changes by single mutations, but it does not prevent single mutations from causing dramatic amino acid changes. We used variation in measurements of the wildtype amino acids at each position (including synonyms) to define a range of WT-like experimental fitness, and assessed the proportion of single and double mutations that were in this range (Figure 3B). Amino acids caused by single mutations were significantly more likely to exhibit WT-like fitness than those caused by double mutations, consistent with the bias of the genetic code for physically conservative amino acids. This difference is significant because of a combination of the quantity and quality of experimental measurements, though the effect size is modest. Our analyses indicate that an amino acid change caused by a double mutation was roughly 80% as likely to display WT-like fitness compared to a change caused by a single mutation.

### Estimated selection coefficients compared to frequency in circulating variants

We compared experimental fitness with the frequency of amino acids at each position in the inhibitor-naïve HIV-1 sequences (Fig. 4). The most frequently observed amino acids in isolates tended to have WT-like fitness in the NL4-3 genetic background under the conditions of our experiments. Among amino acid states at frequency below 0.003 in the isolates, many were null-like in NL4-3. Epistasis provides one explanation for this observation as the fitness of these amino acids could depend strongly on genetic background. In future work, it will be interesting to use the experimental fitness measures to guide analyses of epistasis. For the purposes of this study, we chose to focus on mutations where fitness in our experiments is likely a fair indicator of the fitness or mutational sampling of these mutations in circulating viral populations. Because strongly deleterious alleles are extremely unlikely to rise to high frequency (Ohta 1973), our findings indicate that for commonly (f>0.003) observed amino acids, experiments with NL4-3 provide a reasonable estimate of the fitness effects of mutations in circulating HIV-1. Of note, this does not necessarily mean that epistasis is uncommon or unimportant in HIV-1 protease as epistasis is a general feature of protein evolution (Pollock, Thiltgen et al. 2012) that has been shown to mediate HIV-1 protease function (Hinkley, Martins et al. 2011). Rather, NL4-3 protease is the consensus of clinical sequences, and the common clinical amino acid changes in protease may frequently occur on a genetic background that is similar enough to NL4-3 to have closely related effects.

**Figure 4.**
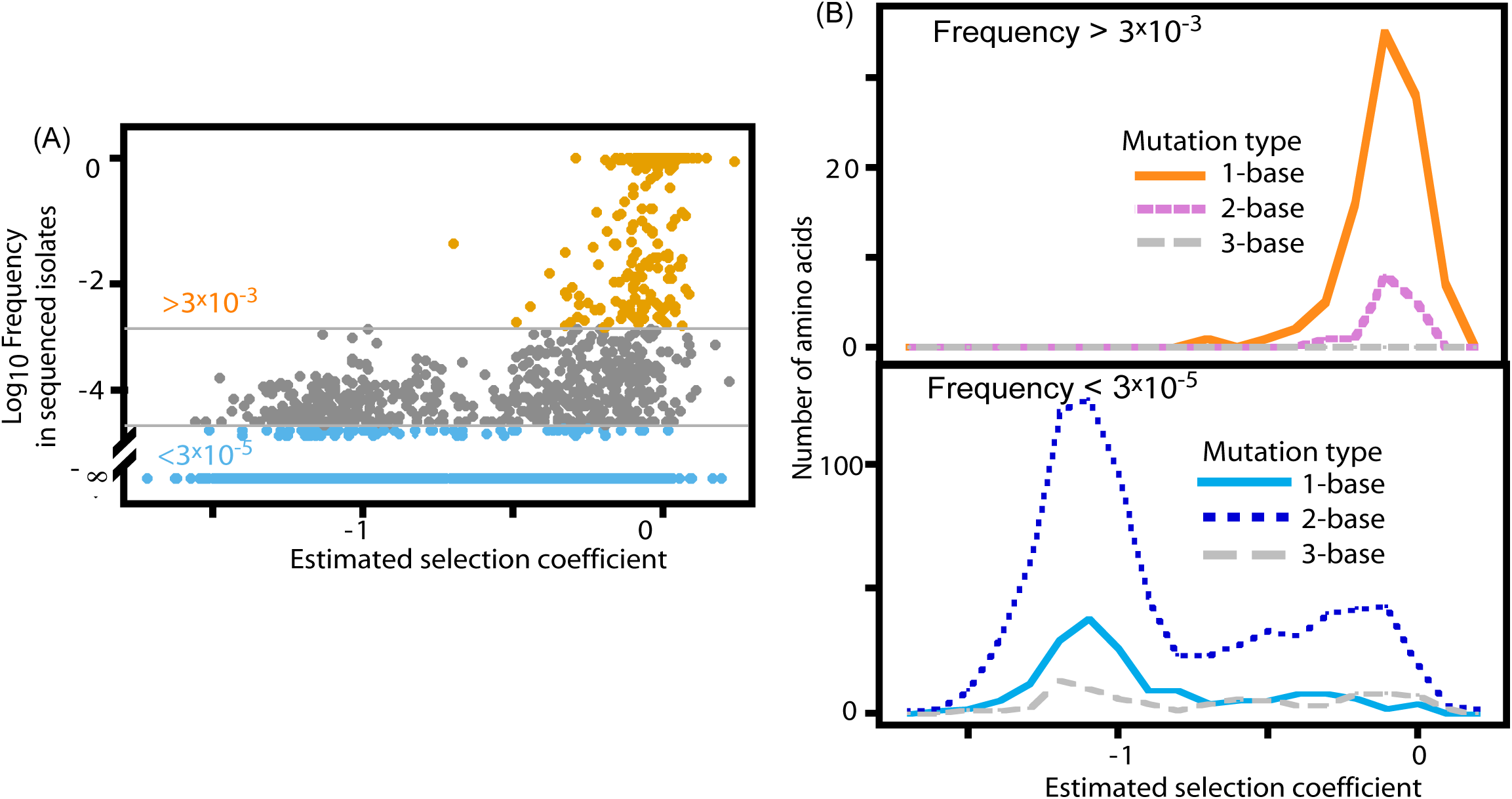
Relationships between experimental fitness and the frequency of mutations in circulating viruses. (A) The frequency of mutations in sequenced viruses from drug-naive patients compared to estimated selection coefficients. Mutations that occurred at a frequency greater than 0.003 are shown in orange, mutations at or below the average frequency of stop codons accessible by single mutations (f<0.00003) are shown in sky blue, and all other frequencies are shown in grey. (B) Distribution of observed selection coefficients for commonly observed (f>0.003) mutations (top panel) and unobserved or very rarely (f<0.00003) observed mutations (bottom panel).

Amino acids that were rarely observed in isolates (grey dots in Fig. 4A) displayed a wide range of experimental fitness in NL4-3. We considered different potential reasons for this range of fitness. We considered the possibility that viruses may expand more readily during infections of human hosts than in cell culture. However this seems unlikely as cell culture is generally a more permissive environment compared to hosts because there is no immune pressure (Vignuzzi, Stone et al. 2006). Epistasis may provide an explanation for the infrequently observed clinical amino acids that have a strong fitness defect in NL4-3, as the genomes where these amino acids occur could have accumulated secondary permissive mutations.

We also considered that null alleles may appear from sequencing errors in the database and/or from sequencing of non-infectious viral particles that could be generated from activated proviruses with genetic defects (Maldarelli, Wu et al. 2014). We cannot distinguish between these mechanisms because they both cause similar appearances of null alleles in isolates. Consistent with these mechanisms, stop codons were present in the protease sequence database. To estimate the likelihood of observing a null protease allele in the database we compared the number of observed stop codons (127) to the number of single-base mutations from the likely ancestral sequence that could lead to a stop codon (98). Based on this ratio we explored how many null amino acids we may expect to find in the sequenced isolates. We defined amino acids as null if they had estimated selection coefficients within two standard deviation of the average stop codon in the EMPIRIC scan (Supplementary Table 1). There were 1001 single mutations that could lead to a null amino acid. Based on the ratio of stop codons to single-base mutational pathways to stop codons, we expected roughly 1300 null amino acids accessible by single nucleotide mutations in the clinical database. We observed 1061, which represents 82% of expectations based on the number of stop codons in the database. These findings indicate that termination codons and most experimentally null alleles seen in the database are likely generated by similar mechanisms, which could be errors in sequencing or non-viable viral lineages.

As each stop mutation accessible by a single mutation from the consensus was observed about once in sequenced isolates, we used this as a cutoff for amino acids that are likely either unfit and/or poorly sampled in natural evolution. Similarly, we used a frequency cutoff >0.003 to delineate mutations that are likely highly fit and well sampled in natural evolution (Fig. 4A). 86% of high frequency amino acids were accessible by single mutations (Fig. 4B), consistent with mutational sampling having a large impact on the genetic diversity of HIV-1 protease. 17% of infrequently observed amino acids were accessible by single mutations. These infrequently observed single mutations were predominantly strongly experimentally deleterious (Fig. 4B), indicating that single mutations that are experimentally fit are also fit and efficiently sampled during the natural evolution of HIV-1. In contrast to single mutations, the experimental fitness distribution of infrequent double mutations was bimodal with peaks that overlapped either null or neutral. The neutral peak indicates that many fit double mutations are inefficiently mutationally sampled during natural evolution. Together, these observations indicate that HIV-1 thoroughly samples single nucleotide mutations, consistent with previous modeling (Perelson 2002); and selection is a primary determinant of infrequent single-base mutations in sequenced isolates.

### Predicting mutation frequency in sequenced isolates

There are multiple factors that complicate comparison of the local protein fitness landscape with amino acid frequency observed in isolates including experimental noise, mutational sampling, and epistasis. Because of noise in our experiments, actual selection coefficients may differ from our estimates. The frequency of mutations in isolates is mediated by a combination of selection and mutational probabilities. Because the protein fitness landscape that we determined is based only on selection, adding a mutational model should improve the prediction of the frequency in isolates. Epistasis has been shown to impact frequencies in sequenced isolates (Flynn, Haldane et al. 2017) as the probability of a mutation depends on fitness in different genetic backgrounds (Hinkley, Martins et al. 2011). Both epistasis and experimental noise will contribute to differences between predicted and actual frequencies.

We considered these complications as we explored relationships between amino acid frequency in sequenced isolates and the experimental protein fitness landscape (Fig. 5). Estimated selection coefficients correlate with amino acid frequency in isolates (Fig. 5A) largely driven by a cluster of mutations with severe experimental fitness defects that were also rarely or never observed in sequenced isolates. This cluster is consistent with the precision of our estimated selection coefficients to clearly distinguish strongly deleterious mutations from those that are capable of efficient replication. In addition, these results indicate that epistasis in protease among drug-naïve hosts infrequently leads to genetic backgrounds that rescue mutations with null-like fitness defects in the likely ancestral background.

**Figure 5.**
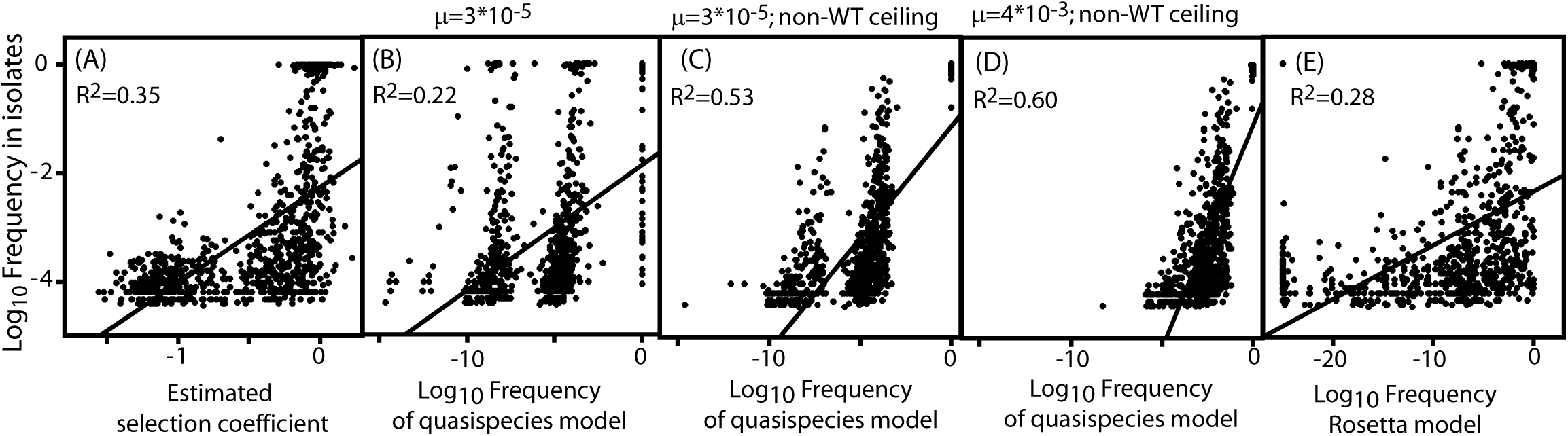
Capability and challenges in predicting amino acid frequency in sequenced isolates. (A) Correlation between amino acid frequency in sequenced isolates and estimated selection coefficients. Unobserved amino acids were omitted from these analyses because because we do not have quantifiable estimates of their likely frequency. (B) Amino acid frequencies were estimated using a quasispecies model based on estimated selection coefficients and the mutation rate of HIV measured in cell culture, (C) An artificial fitness ceiling was applied to non-WT mutations in order to match the dominant quasispecies sequence with wildtype. (D) As in panel C except that the quasispecies model used the error rate estimated from observations of premature stop codons in hosts. (E) Correlation between frequencies predicted based on previously published (Humphris-Narayanan, Akiva et al. 2012) structural modeling of the impacts of mutations on the folding stability, dimerization and binding to substrates of HIV-1 protease.

To investigate contributions of mutational sampling to amino acid frequency, we used a steady-state quasispecies model that combined experimental fitness without epistasis and a simple mutational model based on the error rate (3×10^−5^ per site per generation) during HIV-1 replication in cell culture (Fig 5B). This approach did not improve the correlation between the fitness landscape and frequency in sequenced isolates. Of note, the quasispecies model caused a stratification of amino acids that are visible as four vertical groupings of data points in Fig. 5B. In addition, the quasispecies model predicted rare amino acids that were orders of magnitude lower in frequency than most corresponding observations in isolates. In Fig 5B, the right-most vertical group corresponds to the amino acid at each position with the highest estimated experimental fitness, which resulted in a dominant steady-state sequence that differed at multiple positions from the likely ancestor of the isolates. The apparent adaptive mutations in the fitness landscape are consistent with expected experimental noise (Fig. 2) which has a large impact on predicted frequency for these mutations.

We were concerned that differences between the dominant quasispecies sequence and the likely ancestral sequence of the isolates might complicate further interpretations. Based on the assumption that the likely ancestral sequence was strongly selected and thus highly fit and that estimated adaptive mutations were largely due to measurement error, we explored fitness landscapes that we artificially manipulated by setting a fitness ceiling for non-wildtype amino acids (Fig 5C). We examined different ceilings and found that they resulted in similar correlations between predicted and observed frequency. For presentation purposes we used a ceiling that favored wildtype amino acids by 4% in fitness relative to other amino acids. Adding this fitness ceiling improved the correlation between modeled and isolate frequencies, indicating that noise in experimental measurements around neutral fitness can translate into large impacts on predicted frequency.

Because the use of a fitness ceiling for non-WT amino acids is artificial, we were cautious to limit further analyses to properties such as mutation rate that may be robust to this manipulation. In the quasispecies model, mutation rate will influence the likelihood of single, double, and triple mutations. Since the ceiling is applied to each of these types of amino acid changes, it should not have a large impact on the predicted likelihood of single, double, and triple mutations. Indeed, the second through fourth vertical groups were qualitatively similar to the unadjusted fitness landscape. In both models these groups consisted of amino acids that are 1-, 2-, and 3-mutations away from the sequence with the highest fitness. In the ceiling model, there are fewer apparent triple mutations in isolates because the dominant sequence is the wildtype. With the ceiling, a linear fit (Fig. 5C) still predicts rare amion acids orders of magnitude lower than observations in isolates. Together, these observations indicate that the mutation rate in the model is too low to account for the observed frequencies. We considered how experimental noise and epistasis may influence the observed correlations. We expect that experimental noise would be randomly distributed and would not contribute to the observed grouping of single, double, and triple mutations. Similarly, we expect epistasis to favor some amino acids and disfavor others and not to influence the grouping of single, double, and triple mutations. Thus, cautious interpretation of our results indicates that the mutation rate of HIV-1 in hosts may be higher than the mutation rate in cell culture consistent with a recently reported error rate of 4×10^3^ based on the intrapatient frequency of stop codons (Cuevas, Geller et al. 2015).

We examined how the mutation rate measured within hosts impacted the quasispecies model. With a ceiling-manipulated fitness landscape, we find that the higher mutation rate (Fig. 5D) provides an improved correlation between the quasispecies model and frequencies from isolates. Of note the distribution of the number of mutations per variant from this model was similar to the distribution seen in the sequenced isolates (Fig. S4), indicating that the combination of mutational model and ceiling adjusted fitness landscape are sufficient to recapitulate gross features of sequence diversity in isolates. The higher mutation rate also removed the separation between amino acids with one, two and three mutations, and resulted in a slope between predicted and observed frequencies that was closer to one such that the predicted frequency of rare mutations more closely matched observed frequency. Both of these observations suggest that the higher rate more accurately captures the influence of mutational probabilities on HIV-1 protease evolution.

These analyses highlight some of the challenges that remain for understanding and predicting mutations in circulating HIV-1, particularly estimating small effect mutations with the precision required for accurate evolutionary models, and systematically accounting for epistasis. While these challenges remain the focus of ongoing efforts, the current work indicates that experimental fitness landscapes provide improved predictions of amino acid frequencies compared to estimates from protein-structure based models (Fig. 5E). While there is much we don’t understand in detail about mutational frequencies in the natural evolution of HIV-1, including epistasis, experimental protein fitness landscapes are an important benchmark for testing and refining models.

### Mutational walks

For multiple nucleotide mutations, there were many experimentally fit amino acids that were infrequently observed in sequenced isolates (Fig. 4B). Mutational probabilities that limit access to two- and three-base mutations provide a compelling rationale for these observations. According to well established population genetic theory (Kimura 1983), the probability of a neutral mutation is proportional to the rate of the mutation. As single-nucleotide changes are the most common mutation in HIV-1 (Mansky 1996), amino acid changes requiring multiple-base changes should often arise by the serial accumulation of individual independent mutations. The likelihood of this mechanism of multiple-base change is the product of the probabilities of each individual mutation and is less likely than the single-base mutations. The lower likelihood of multiple-base mutations compared to single-base mutations should limit their frequency in populations of HIV-1.

Because simultaneous replacement of two or more nucleotides in a codon is rare (Mansky 1996), mutational walks of separately occurring single-base mutations may provide relevant access to these types of amino acid changes. The likelihood of a mutational walk depends on the fitness effects of intermediate steps (Gillespie 1983; Elena and Lenski 2003). If an intermediate step is strongly deleterious, then the pathway will be unlikely to occur even if the final state is neutral. We examined the intermediates for two-nucleotide mutations (Fig. 6) where we identified a set of both common and unobserved clinical amino acids with similar high fitness. Among the two-base mutations common in circulating variants (Fig. 6A), estimated selection coefficients for intermediate one-base mutations exhibited a narrow and highly-fit distribution. Intermediates for a random sample of equally fit but unobserved two-base mutations (Fig. 6B) exhibited a broader range of effects with some fitness measurements close to null. The fitness of intermediates for these unobserved mutations was significantly lower (p=0.0003) than for intermediates for common circulating mutations consistent with mutational walks contributing to multiple-nucleotide codon changes during natural evolution.

**Figure 6.**
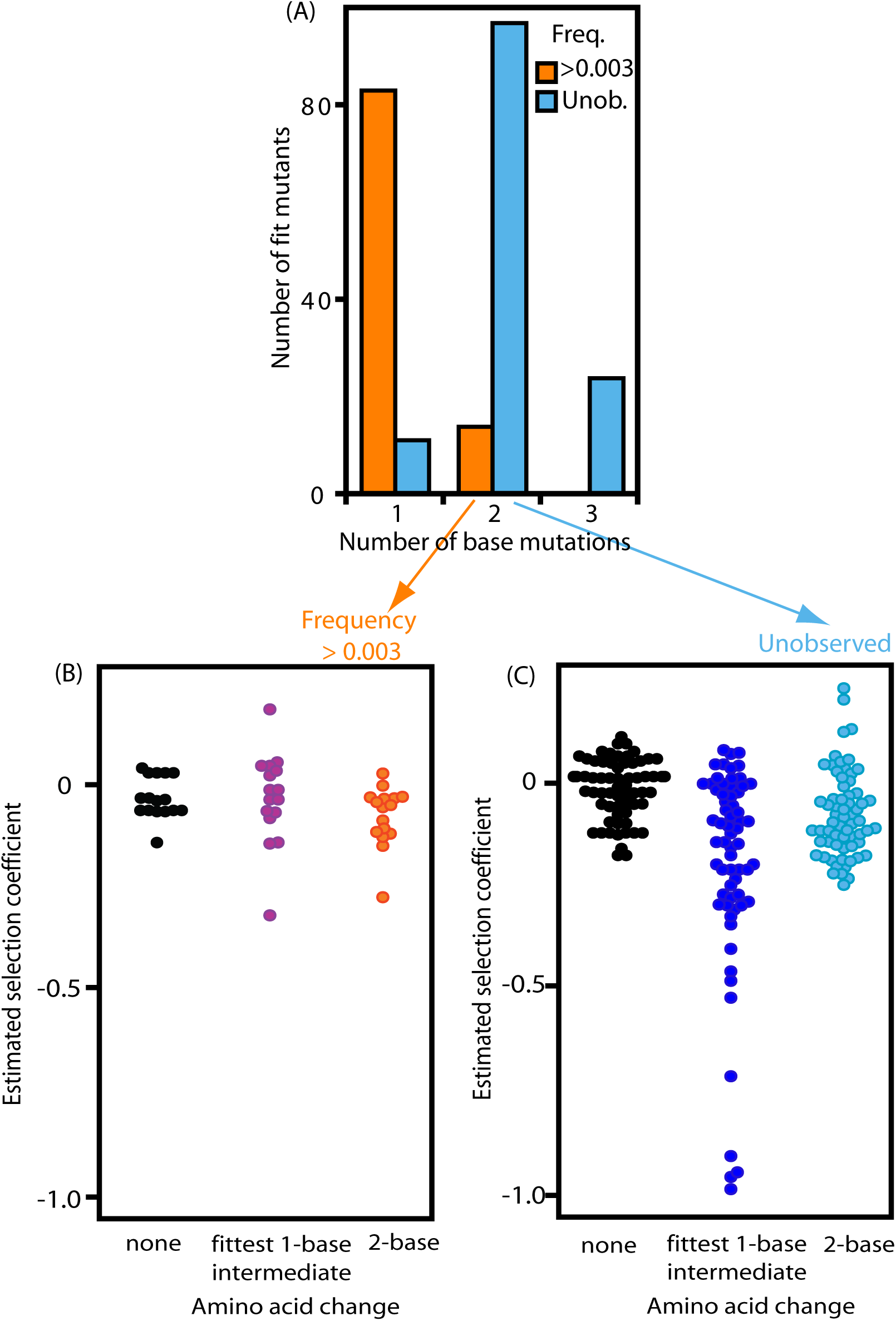
Frequently observed amino acid changes requiring two nucleotide changes are accessible by fit intermediates compared to highly fit amino acid changes that were unobserved. (A) Number of amino acids with experimental fitness similar to commonly observed natural mutations. Amino acids were considered fit if they exhibited selection coefficients were greater than the average minus one standard deviation of commonly observed (frequency > 0.003) mutations. The partitioning of fit 1- and 2-base mutations are statistically different (p=10^−16^) based on a Pearson’s chi-squared test with Yates’ continuity correction. (B) Orange circles show estimated selection coefficients for amino acid changes requiring two nucleotide changes that were observed with a frequency greater than 0.003 in clinical samples. Purple circles show the most fit amino acid intermediate and black circles show on-pathway synonymous codons of the likely ancestral amino acid. (C) Sky blue circles represent a randomly selected set of highly fit amino acid changes requiring two nucleotide mutations that were not observed in the sequenced isolate data set. Blue circles represent the most fit amino acid intermediate and black circles show on-pathway synonymous codons of the likely ancestral amino acid. Based on a Wilcoxon rank-sum test, the intermediates for the frequently observed amino acid changes exhibit higher fitness (p<0.0003) than the intermediates for the unobserved amino acid changes.

### Mutational tolerance

Because mutational pathways likely contribute to the sampling of amino acid changes in circulating variants, we examined if multiple mutations preferentially occurred at amino acid positions that were tolerant to amino acid changes (Fig. 7). Positions in HIV-1 protease tended to be either highly sensitive where most amino acid changes caused null-like phenotypes, or highly tolerant where most amino acid changes caused minor alterations to fitness. If mutational walks contribute to multiple-base mutations, they may be more likely to occur at tolerant positions that would have more opportunities for intermediate steps with high fitness. We first examined amino acid changes in sequenced isolates that were accessible by single mutations. Compared to all positions, we observed single-mutations less frequent at highly sensitive positions, consistent with the idea that positions where most mutations are strongly deleterious are unlikely to accumulate mutations.

**Figure 7.**
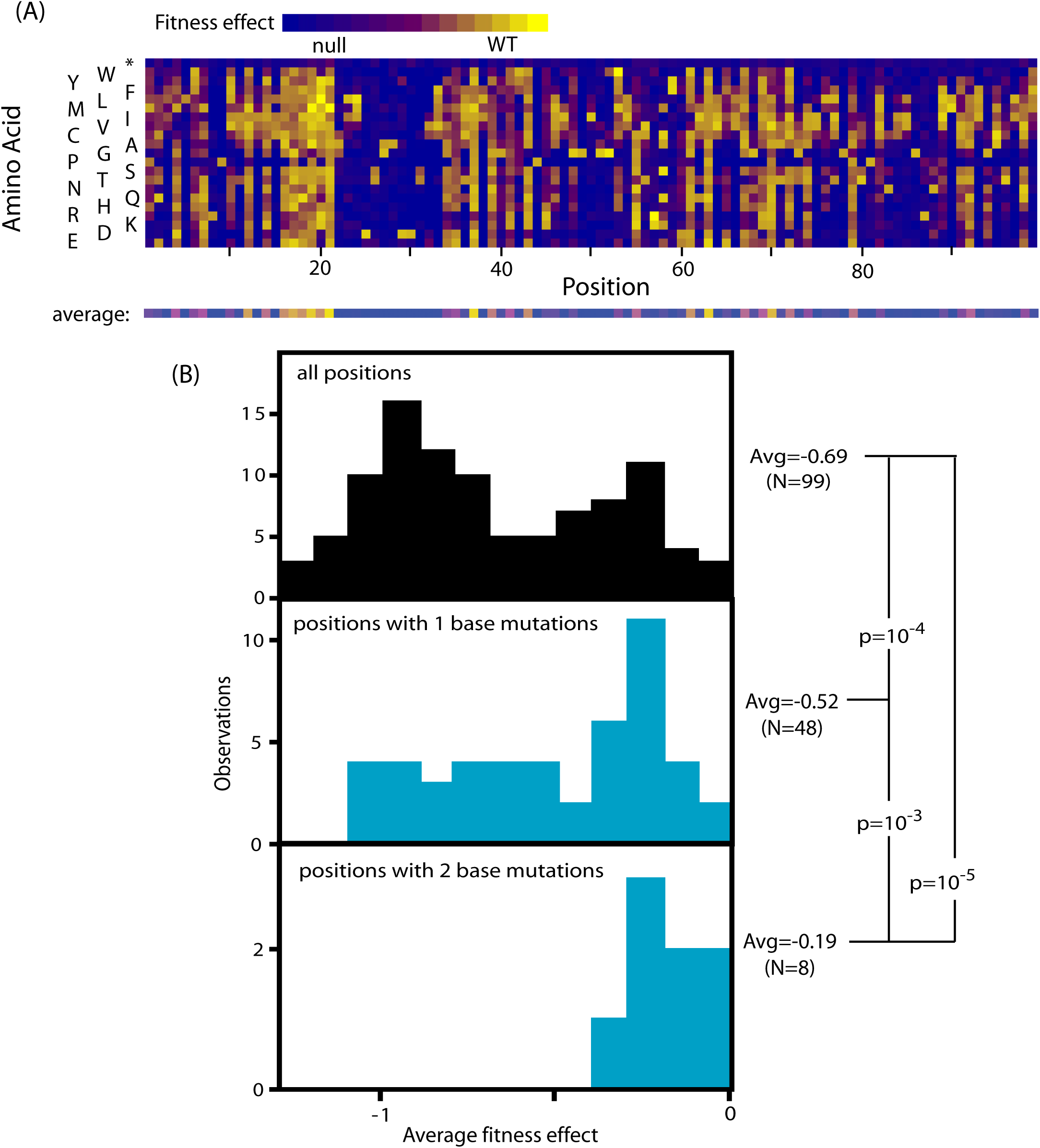
Frequently observed mutations preferentially occur at positions in protease that are relatively tolerant to mutations. (A) Heatmap representation of the estimated selection coefficient of amino acid changes in protease from NL4-3 HIV-1. The average effect of all amino acid changes is shown on the bottom line and these values were used as a measure of the tolerance of a position to mutation in panel B. (B) The overall distribution of mutational tolerance (top panel) is bi-modal with a cluster of positions where the average fitness effect exhibited mild to no fitness effect (s≈0) and a cluster of positions where the average fitness is close to null (s=-1). Positions where mutations were observed in circulating viruses at frequencies above 0.003 were skewed towards higher tolerance (middle two panels). Statistical significance was tested using a one-tailed bootstrap analyses.

Multiple-base mutations in sequenced isolates occurred predominantly at highly tolerant amino acid positions compared to both the overall distribution of sensitivity and the distribution of single mutations (Fig. 7). Because selection is weaker at tolerant positions, they should more freely accumulate mutations during HIV-1 evolution. Such broad peaks in local fitness landscapes provide a greater opportunity for mutational walks to amino acids that involve multiple-base changes.

## Conclusions

Combined analyses of the sequenced isolates and an experimental protein fitness landscape of HIV-1 protease indicate that sampling of multiple-base substitutions in the same codon is limited during HIV-1 evolution. Largely for this reason, the distribution of amino acid changes in circulating variants is skewed towards amino acid changes accessible by single-nucleotide mutations. The mutations that HIV-1 accumulates during genome copying are predominantly single-nucleotide changes that are unlikely to simultaneously occur in the same codon. Therefore, the likelihood of observing multiple mutations depends on the fitness of single mutation intermediates. Because the majority of amino acid changes require multiple mutations, this mechanism of mutational sampling can have a large influence on protein sequence evolution, even for viruses such as HIV-1 that have high genetic diversity in hosts.

Strong evidence indicates that the genetic code was selected to favor conservative amino acid changes by single-nucleotide mutations (Sengupta and Higgs 2015); yet, we observe many multiple-base mutations that support efficient HIV-1 expansion in our experiments. These two observations are not mutually exclusive. Our observations of multiple-base mutations with small fitness effects is at least in part due to a common feature of proteins, the tendency for many sites to be highly tolerant to amino acid changes even for proteins whose sequences are highly conserved in nature (Roscoe, Thayer et al. 2013; Mishra, Flynn et al. 2016). Tolerant sites often permit any amino acid change such that multiple-base mutations at these positions will not exhibit strong defects. Because tolerant positions appear to be a general feature of proteins, our observations that mutational sampling constrains the amino acid sampling of HIV-1 likely extend to many other proteins and organisms.

## Methods

### Library construction

To facilitate the initial introduction of mutations, protease plus 50 bases of upstream and downstream flanking sequence bracketed by KpnI sites was cloned from pNL4-3 into pRNDM (Hietpas, Roscoe et al. 2012). Each codon of protease in the pRNDM plasmid was individually subjected to site saturation mutagenesis using a cassette ligation strategy (Hietpas, Roscoe et al. 2012). A pNL4-3Δprotease plasmid was generated to efficiently accept protease variants from the pRNDM construct. The pNL4-3Δprotease plasmid was constructed with a unique AatII restriction site. The pNL4-3Δprotease plasmid was treated with AatII enzyme followed by T4 DNA polymerase without nucleotides to remove the 3’ overhang. Protease variant libraries in pRNDM were excised with KpnI and treated with T4 DNA polymerase without nucleotides in order to remove 3’ overhangs. The protease variant libraries and treated pNL4-3Δprotease samples contained 25 bases of complementarity at both ends and this facilitated efficient assembly using Gibson Assembly. All enzymes were from New England Biolabs. Samples with mutations at 9-10 consecutive positions in protease were pooled to generate 10 library samples that together include mutations at each of the 99 positions in protease.

### Growth competition

Viral recovery and competitions were performed similar to previous descriptions (Duenas-Decamp, Jiang et al. 2016). Briefly, 2.5 μg of plasmid DNA encoding full length HIV-1_NL4-3_ was transfected into 293T cells using calcium phosphate. Supernatant of recovered P0 viral libraries was harvested after 48 hrs, clarified by filtration through 0.45 μm filters and stored at −20°C. We used an RT assay, which quantifies reverse transcriptase activity by real-time PCR, to normalize virion production (Vermeire, Naessens et al. 2012). Viral infections, including technical replicates, were performed using 5.0×10^8^ RT units of virus (P0) and 3.0×10^6^ Jurkat T cells in 500 μL RPMI complete media for 2 hours. Cells were washed twice with sterile PBS and seeded in 1.5 mL of RPMI complete media in a 24-well plate. P1 viral supernatant was collected and cells were split on days 2, 4, 8, 11, 14, and 16. Fresh media was replaced after each collection time-point. Viral RNA was isolated from P1 viral supernatant collected on days 4 and 8 using Qiagen’s QIAamp MinElute Virus Spin kit. Ultracentrifugation of 300 μL of viral supernatant was performed to pellet virions using a Beckman Optima Max XP ultracentrifuge with a TLA-55 rotor at 25,000 rpm for 1 hour at 4 °C. Supernatant was removed and virion-associated RNA in the pellet was isolated according to the kit protocol, and eluted in 20 μL.

### DNA preparation and sequencing

Samples were prepared for sequencing essentially as previously described (Duenas-Decamp, Jiang et al. 2016). HIV-1 genomic RNA was extracted from supernatants containing virions using High Pure Viral RNA kit (Roche Inc.). SuperScript III and a primer downstream of the randomized regions were used to reverse transcribe viral RNA to cDNA. The cDNA samples were processed as previously described (Hietpas, Roscoe et al. 2012) to add barcodes to distinguish pre- and post-selection samples as well as to add base sequences needed for Illumina sequencing. In our previous work we optimized and tested the accuracy of the sequence based readout of variant frequency. Importantly, base call errors from processing and sequencing were a small fraction of variant frequency in the initial libraries for the vast majority of mutations using this procedure (Duenas-Decamp, Jiang et al. 2016). Sequencing data from the current work has been submitted to the Short Read Archive under BioProject ID PRJNA476312.

### Sequence analysis

Viral RNA samples drawn from transfected cells at days 0 (initial transfection from HEK293 cells to T-cells), 4 and 8 were processed using Illumina 36-bp single read sequencing on a Genome Analyzer II. Reads with a Phred score of 20 or above (>99% confidence) across all 36 bases were used for time-dependent analysis. Using these counts, for each species, *i*, the selection-rate constant (Lenski, Rose et al. 1991), relative to the mean Malthusian fitness of stop codons can be defined as the slope of a normalized logarithmic abundance versus time

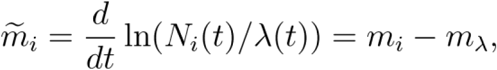

where *N*_*i*_(*t*) refers to counts at time *t*, *m*_*i*_ is the un-normalized Malthusian fitness, and *λ*(*t*) are normalization counts that correspond to the Malthusian fitness of stop codons such that

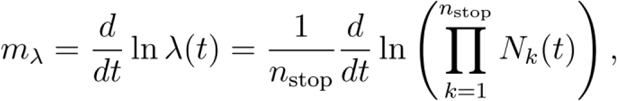

where *n*_*stop*_ is the number of stop codons in the library. This normalization convention ensures that the mean fitness of stop codons is zero. We estimated the relative Wrightian fitness of each variant by normalizing to the Malthusian fitness of the wildtype sequence 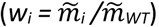, such that *w*_*WT*_ = *1*. Selection coefficients were calculated as *s*_*i*_ = *w*_*i*_ − *1*. In order to collect measurements across the entire protease sequence, a set of 10 non-overlapping EMPIRIC libraries was prepared, each one spanning 9 or 10 contiguous amino acid positions. For three of these libraries, biological replicate experiments were performed and analyzed using the same approach.

### Analyses of protease sequences from circulating isolates

Sequence information for HIV-1 protease from circulating isolates was collected from the Stanford University HIV drug resistance database (Rhee, Gonzales et al. 2003; Shafer 2006) in April, 2017 using the Genotype-Rx protease data-set. The protease sequences were restricted to include only subtype B strains that were annotated “None” for treatment with protease inhibitors and had a complete nucleotide sequence (i.e. “NASeq”). These requirements resulted in 32,163 sequences from 30,987 unique subjects. 11,487 sequences contained at least one position with an annotated mixture of amino acids (e.g. “KR” represents a mixture of lysine and arginine). At positions annotated with mixtures, each amino acid in the reported mixture was counted as an equal fraction of the mixture. When multiple sequences were present from the same individual, we weighed each sequence by the reciprocal of the number of sequences, such that every host was equally represented. We did not count stop codons because they represent known null alleles, of which there were 127 in the data set. We examined the impacts of excluding mixtures on the relationship between amino acid frequency and fitness (Fig. S5). Excluding sequences that contained mixtures did not dramatically alter the correspondence between frequency and experimental fitness. We examined the location of amino acid mixtures (Fig. S6) and note that they tend to occur at positions that exhibit the most amino acid divergence between different isolates. This observation is consistent with many of the mixtures representing intra-host variation and was a motivation for including reported amino acid mixtures in our analyses.

### Modeled populations

While data collected via EMPIRIC provide a direct readout of relative fitness for the various species in the library, under certain population dynamics assumptions these fitness measurements can also be used to extrapolate initial (constructed) populations to those at later times.

The quasispecies model (Eigen 1971; Eigen, McCaskill et al. 1988) has previously been used to describe the evolutionary dynamics of HIV-1 (Nowak, May et al. 1990). Since our measurements were done on libraries of single codon substitutions that did not include higher order substitutions, we consider an additive quasispecies model where the codon and amino acid frequencies at different positions are independent of one another. The discrete-time quasispecies model for the codon frequencies is

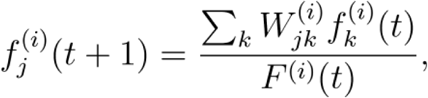

where 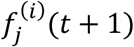 is the frequency of codon *j* at position *i* at time t+1, 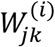 is the mutation-selection matrix describing the fitness change caused by mutation of codon *j* to codon *k* at position *i*, 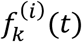 is the frequency of codon *k* at position *i* at time *t*, and *F*^(*i*)^(*t*) is the average fitness of the population at time *t* and is calculated as,

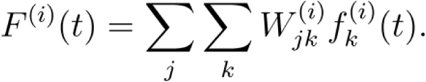

The mutation selection matrix, 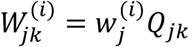 is the product of a site-specific fitness landscape, 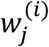 defined by the EMPIRIC data and a single-site mutational landscape (Jain and Krug 2007),

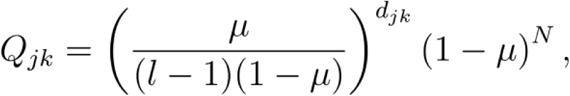

where each residue position implies a (single codon) sequence length *N=3*, and *l=4* represents the possible nucleotides at any position in this sequence, *μ* is the mutation rate and *d*_*jk*_ is the Hamming distance between codons *j* and *k*.

We computed the steady state solution to the quasispecies model, given by the dominant eigenvector of 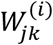. In order to predict steady state frequency distributions under this model, two different mutation rates were employed (see Fig. 5): an error rate of 3 × 10^−5^ previously measured in cell culture (Mansky 1996), and an error rate of 4 × 10^−3^ based on observations of premature stop codons observed in hosts (Cuevas, Geller et al. 2015). When the fitness used in our quasispecies model included a ceiling that favored wildtype amino acids by 4%, the higher of these two mutation rates led to an accurate description of the mean number of mutations in the steady state population (Fig. S4).

## Acknowledgements

We are thankful to L. Jiang for guidance and helpful advice related to the EMPIRIC experiments. We are also thankful to Tanya Kortemme for sharing free energy data from structural modeling. This work was supported by grant P01GM109767 from the National Institutes of Health.

